# Characterisation and genomic analysis of bacterial nutritional endosymbionts in Australian ticks from shotgun metagenomic sequencing

**DOI:** 10.64898/2026.08.12.744556

**Authors:** Laurene Leclerc, Julia Meltzer, Xabier Vázquez-Campos, Olivier Duron, Julien Amoros, Brendan P. Burns, Nathan Lo

## Abstract

Ticks are obligate hematophagous arthropods and feed exclusively on blood. As blood is nutrient-poor, ticks rely on bacterial endosymbionts to synthesise nutrients, yet the diversity and functional roles of these symbionts in Australian ticks remain largely uncharacterised. This is critical to address as these ticks are of high medical importance in Australia. In this study, shotgun metagenomic sequencing was performed on *Bothriocroton concolor*, *Bothriocroton hydrosauri*, *Haemaphysalis longicornis* and *Ixodes holocyclus*, enabling the recovery of six complete or partial metagenome-assembled genomes (MAGs). These comprised *Coxiella*-like endosymbionts (CLE), a facultative *Rickettsia* symbiont, and two *Midichloria mitochondrii* strains (Ixholo1 and Ixholo2). Functional annotation of these taxon-specific symbionts revealed the absence of virulence factors and the presence of B-vitamin and/or heme biosynthesis genes, indicative of nutritional mutualism, which is essential for tick hematophagy. The CLEs additionally harbour genes of the shikimate pathway, which modulate blood feeding in ticks by regulating serotonin biosynthesis. Furthermore, functional annotation and pangenomic analysis of *Midichloria* spp. found evidence that the genus may encompass multiple species, as well as the retention of genes potentially associated with an intramitochondrial lifestyle in *M. mitochondrii* Ixholo2. Tick microbiomes are dominated by non-pathogenic microorganisms, which are often overshadowed by pathogens. These include the endosymbionts, which can influence host biology and pathogen transmission, and are fundamental for the development of diagnostic tools and taxon-specific tick biocontrols.

## INTRODUCTION

Ticks are the second most important vector after mosquitoes, transmitting bacterial, viral, nematode and protozoan pathogens to humans and animals worldwide (1–4). Novel tick-borne pathogens are continuously being discovered and the global burden of already described pathogens is increasing (5–10). In addition to pathogens, ticks harbour bacterial endosymbionts. These endosymbionts influence tick nutrition, fitness, reproduction, immunity and disease ecology by facilitating or competing with pathogens (11). Ticks are obligate hematophagous arthropods and feed exclusively on blood. As blood is nutrient-poor, ticks have evolved mutualistic interactions with obligate bacterial symbionts to synthesise nutrients, specifically B vitamins such as biotin, folate and riboflavin (12–14). Most ticks typically harbour a single lineage of nutritional symbiont, such as a *Coxiella*-like endosymbiont (CLE), *Francisella*-like endosymbiont (FLE), *Rickettsia* endosymbiont (RE) or *Candidatus* Midichloria mitochondrii (hereafter *M. mitochondrii*) (12–14). Dual endosymbiosis has also been described in the tick *Hyalomma marginatum*, where riboflavin is synthesised by a FLE, biotin by *M. mitochondrii*, and folate and heme by both (15). Ticks also harbour facultative symbionts that are not essential for host survival but may nevertheless influence tick physiology, ecology and vector competence (16). Some facultative symbionts, particularly certain members of the genera *Rickettsia* and *Rickettsiella*, harbour *cif* genes encoding cytoplasmic incompatibility factors and may manipulate tick reproduction, similarly to *Wolbachia* in other arthropods (17, 18).

At least 18 of the 73 Australian tick species have been shown to feed on humans and domestic animals (19). The tick with the highest medical importance in Australia is *Ixodes holocyclus*. Bites from this species can cause paralysis through the transmission of toxins, mammalian meat allergies (resulting from an IgE-mediated hypersensitivity reaction to the oligosaccharide galactose-α-1,3-galactose), as well as infectious diseases caused by pathogen transmission. *Ixodes holocyclus* is also known to harbour two strains of the endosymbiont *M. mitochondrii,* Ixholo1 and Ixholo2 (20, 21). Genomic studies of Ixholo1 indicate it synthesises heme, biotin and folate for its host, and is involved in oxidative stress response and osmotic regulation (21, 22). Research on Ixholo2, as well as other Australian tick endosymbionts, is lacking.

To characterise symbionts and, more broadly, the tick microbiome, previous studies have commonly used 16S rRNA gene amplicon sequencing and PCR (23–33). These studies have provided evidence of novel species from genera harbouring tick-borne pathogens and symbionts, including *Coxiella, Ehrlichia, Francisella, Neoehrlichia, Rickettsia, Anaplasma* and *Borrelia* species (24, 25, 29, 32, 33). Amplicon sequencing is useful for providing a snapshot of bacterial communities (34, 35). Other studies have also used RNA sequencing to characterise the tick microbiome (26, 36, 37). While this method is useful for the detection of RNA viruses and providing functional insight into the microbiome, it only sequences a subset of the total genetic material and does not sequence bacterial genomes.

Shotgun metagenomics is a culture-independent method that allows the investigation of complex microbial communities by characterising and functionally annotating microbial genomes (38). This method can be optimised to ensure high sequencing depth to construct metagenome-assembled genomes (MAGs), allowing the diversity, abundance, and functional potential of these microorganisms to be determined. Important genes and pathways can therefore be characterised, including those involved in symbiosis and pathogenicity. While tick shotgun metagenomics is becoming increasingly common around the world, only one study has applied this method to Australian ticks, namely *Ixodes tasmani*, *I. holocyclus* and *Haemaphysalis bancrofti* (39). This study characterised and sequenced MAGs of five novel DNA viruses (four Anelloviruses and a Circovirus) and a novel SFG *Rickettsia* isolate (39). Little is known about the metagenomes of other Australian tick species, highlighting the need for further community-wide sequencing. We therefore aimed to use shotgun metagenomic sequencing on four species of Australian ticks to: 1) sequence and annotate endosymbiont MAGs, 2) perform phylogenetic analyses of the MAGs, 3) characterise genes and pathways involved in symbiosis to understand the symbiotic relationships of the bacteria and their tick host, and 4) perform a pangenomic analysis of *Midichloria* spp. to postulate the role and intramitochondriality of *M. mitochondrii* Ixholo2.

## METHODS

### Tick collection

Questing adult female and nymphal *I. holocyclus* were collected in November 2023 by dragging a 1 m^2^ white cloth over vegetation in Chowder Bay, Sydney, Australia. The ticks were kept at ambient temperature and dissected live. All other ticks were collected by citizen scientists throughout Australia between 2017 and 2024. Specimens were examined utilising Leica M60 and M250 C stereo microscopes, and stored long-term at −30 °C. The species, sex and life stage for each tick were determined using taxonomic keys (19).

### Tick preparation, DNA extraction and DNA sequencing for shotgun metagenomic sequencing

Microbial and tick DNA was extracted for metagenomic sequencing from *Bothriocroton concolor* (n=1), *Bothriocroton hydrosauri* (n=1), *Haemaphysalis longicornis* (n=2) and *Ixodes holocyclus* (n=16). Tick whole bodies were washed in 96% ethanol by vortexing for 60 s, followed by a 30 s wash in phosphate-buffered saline (PBS). The abdominal contents of adult ticks were dissected and stored in 180 μL of PBS at −80 °C. Nymphal ticks were cut medially using a scalpel to enhance homogenisation. All samples were homogenised using a micro pestle. The homogenate was centrifuged at 17,000 x g and the supernatant was filtered through polyethersulfone (PES) filters (0.45 µm or 0.8 µm) (Sartorius, Germany) for microbial enrichment (Table 1). DNA extraction was performed using the Qiagen DNeasy Blood & Tissue Kit (QIAGEN, Valencia, California, USA) according to the manufacturer’s protocol. To improve digestion, samples were incubated in buffer AL or buffer ATL for 1 hour to overnight (Table 1). For all extractions, the quantity and purity of DNA was determined by Qubit 4 Fluorometer (Thermo Fisher Scientific, USA). The DNA samples were stored at −30 °C until further use. To validate our DNA extraction and bacterial enrichment protocols, five samples were subjected to 16S rRNA gene (V1-3) amplicon sequencing, which was performed on extracted DNA at the Ramaciotti Centre for Genomics on an Illumina MiSeq v3 2×300 base pairs (bp) sequencing run using PCR primers 27f and 519r (Supplementary Methods, Supplementary Figure 1, 2 and 3, Supplementary Tables 1 and 2). Following validation, Illumina DNA library preparation and DNA sequencing (NovaSeq 6000 SP 2×150 bp flowcell) were performed on the 20 tick DNA samples by the Australian Genome Research Facility (AGRF) and produced between 22 and 38 million reads per sample (Table 1).

**Table 1.** Tick samples used for shotgun metagenomic sequencing, with corresponding microbial enrichment and cell lysis methods.

| ID | Tick species | Life stage, sex | Engorgement (host if known) | Collection location | Microbial enrichment | Lysis buffers and enzymes for digestion | Data yield (bp) |
| --- | --- | --- | --- | --- | --- | --- | --- |
| 4BC1 | *Bothriocroton concolor* | Adult female | Partially fed (echidna) | Unknown | 0.8 µm PES filter | 3h digest with buffer AL & proteinase K | 8.27 Gb |
| 5BH2 | *Bothriocroton hydrosauri* | Adult female | Partially fed (lizard) | Cape Leeuwin, WA (Jul 2024) | No enrichment | 3h digest with buffer AL & proteinase K | 9.23 Gb |
| 8HL1 | *Haemaphysalis longicornis* | Adult female | unfed | Lake Haven, NSW (Oct 2017) | 0.8 µm PES filter | 3h digest with buffer AL & proteinase K | 9.75 Gb |
| 9HL2 | *Haemaphysalis longicornis* | Adult female | unfed | Lake Haven, NSW (Oct 2017) | No enrichment | 3h digest with buffer AL & proteinase K | 8.78 Gb |
| 11A2 | *Ixodes holocyclus* | Adult female | engorged | Unknown | 0.45 µm PES filter | 1h digest with buffer AL & proteinase K | 10.18 Gb |
| 12A3 | *Ixodes holocyclus* | Adult female | engorged | Unknown | 0.8 µm PES filter | 1h digest with buffer AL & proteinase K | 8.24 Gb |
| 13A4 | *Ixodes holocyclus* | Adult female | engorged | Unknown | 0.8 µm PES filter | 1h digest with buffer AL & proteinase K | 7.97 Gb |
| 14A5 | *Ixodes holocyclus* | Adult female | engorged | Unknown | 0.45 µm PES filter | 1h digest with buffer AL & proteinase K | 9.96 Gb |
| 15A6 | *Ixodes holocyclus* (pool of 5) | Nymph | unfed | Sydney, NSW (Nov 2023) | 0.45 µm PES filter | 2h digest with buffer AL & proteinase K | 9.53 Gb |
| 16IH4 | *Ixodes holocyclus* (pool of 3) | Nymph | unfed | Sydney, NSW (Nov 2023) | No enrichment | O/N digest with buffer ATL & proteinase K | 11.48 Gb |
| 17IH1 | *Ixodes holocyclus* | Adult Female | unfed | Sydney, NSW (Nov 2023) | 0.8 µm PES filter | 3h digest with buffer AL & proteinase K | 10.18 Gb |
| 18IH2 | *Ixodes holocyclus* | Adult Female | unfed | Sydney, NSW (Nov 2023) | No enrichment | 3h digest with buffer AL & proteinase K | 9.74 Gb |
| 19A14 | *Ixodes holocyclus* | Adult female | Unfed | Sydney, NSW (Nov 2023) | No enrichment | 3h digest with buffer AL & proteinase K | 10.39 Gb |
| 20IH5 | *Ixodes holocyclus* | Adult female | Unfed | Sydney, NSW (Nov 2023) | No enrichment | 3h digest with buffer AL & proteinase K | 8.5 Gb |

### De novo metagenomic assembly pipeline

Raw reads were trimmed to remove adapter and low-quality sequences, and separated into forward reads, reverse reads and singletons using BBmap (v.38.63) (40). Libraries were quality-checked using FastQC (v0.11.9) (41) and then concatenated by species. The raw reads were *de novo* co-assembled by tick species using SPAdes (--phred-offset 33, -k 21,33,55,77,99,111,127) (v3.15.5) (42, 43). As the *I. holocyclus* co-assembly was too large for SPAdes, host reads were removed. This was done by assembling the reads using MEGAHIT (v1.2.9) (44) and identifying Eukaryotic contigs using Whokaryote (v1.1.2) (45). The trimmed reads were then mapped to the Eukaryotic contigs using Bowtie2 (v2.5.2) (46), and all unmapped reads were extracted using Samtools (v1.20) (47) and reassembled using SPAdes (v3.15.5) (42, 43) (--phred-offset 33, -k 21,33,55,77,99,111,127). Sample-wise abundance was then estimated by mapping reads to the contigs using Strobealign (v. 0.16.1) (48) with the --aemb flag, and the taxonomy of each contig was determined using MMseqs2 (v18-8cc5c) (49). The Strobealign abundance and MMseqs2 taxonomy were then used by TaxVamb (v5.0.5) (50) to bin the contigs. The contig abundance and taxonomic data were then also used to analyse the bacterial composition of each library, which was plotted in a histogram using ggplot2 (v3.5.2) (51) and Plotly (v4.11.0) (52) in RStudio (v4.4.1) (53) (Supplementary Figure 4). The MAG completeness and contamination were determined using CheckM2 (v1.0.2) (54) and BUSCO (v5.8.2) (Table 2) (55). Then, the taxonomic classification of the bacterial genomes was performed using GTDB-Tk (v2.5.2) with GTDB r226 (56), and the pairwise identity of the MAG against its closest genome reference (from GTDB-Tk) was calculated using MAUVE (v1.1.3) (57) using the Mauve Contig Mover algorithm (Table 2) (Supplementary table 3). Finally, the bacterial MAGs were annotated with Prokka (v1.14.6) (58) and genes involved in endosymbiosis, including heme, biotin, folate and riboflavin biosynthesis, were identified in the annotated genomes. Homologs of the *cif* genes were searched for using Orthofinder analyses, as described in Amoros et al. (2025) (18).

**Table 2.** Genomic features of MAGs. BUSCO Score abbreviations: C = 254 Complete, S = Complete and single-copy, D = Complete and duplicated, F = Fragmented, M = 255 Missing.

|  | *Coxiella*-like endosymbiont of *Haemaphysalis longicornis* | *Midichloria mitochondrii* strain Ixholo1 | *Midichloria mitochondrii* strain Ixholo2 | *Coxiella*-like endosymbiont of *Bothriocroton hydrosauri* | *Coxiella*-like endosymbiont of *Bothriocroton Concolor* | *Rickettsia* endosymbiont of *Bothriocroton concolor* |
| --- | --- | --- | --- | --- | --- | --- |
| Tick host | *Haemaphysalis longicornis* | *Ixodes holocyclus* | *Ixodes holocyclus* | *Bothriocroton hydrosauri* | *Bothriocroton concolor* | *Bothriocroton concolor* |
| Assembly level | Complete | Complete | Partial | Partial | Partial | Partial |
| CheckM2 completeness | 99.97% | 96.29% | 71.5% | 66.82% | 51.18 % | 49.22% |
| CheckM2 contamination | 0.01% | 0% | 0.11% | 2.54% | 0.53% | 1.0% |
| BUSCO Score | C:90.5%<br>[S:90.5%,D:0.0%],<br>F:3.4%,M:6.0% | C:85.3%<br>[S:85.3%,D:0.0%],<br>F:3.4%,M:11.2% | C:55.2%<br>[S:55.2%,D:0.0%],<br>F:1.7%,M:43.1% | C:55.2%<br>[S:55.2%,D:0.0%],<br>F:17.2%,M:27.6% | C:31.9%<br>[S:31.9%,D:0.0%],<br>F:17.2%,M:50.9% | C:44.8%<br>[S:44.8%,D:0.0%],<br>F:5.2%,M:50.0% |
| Genome size | 997,623 | 937,896 | 812,790 | 932,107 | 953,516 | 535,292 |
| GC content | 0.38 | 0.32 | 0.35 | 0.39 | 0.41 | 0.32 |
| Scaffolds | 1 | 1 | 80 | 241 | 220 | 149 |
| N50 | 997,623 | 937,896 | 13,241 | 4,155 | 4,784 | 3,680 |
| Total coding sequences | 793 | 1,110 | 874 | 1,014 | 1,263 | 638 |
| GTDB-Tk Closest relative | *Coxiella*-like endosymbiont strain 580 chromosome (GCF_024730665.1) | *Candidatus* *Midichloria* sp. isolate IholoSidney (GCA_030060845.1) | (GCF_000219355.1) *Candidatus* *Midichloria mitochondrii* IricVA | N/A | N/A | N/A |
| % pairwise identity to closest relative with Mauve alignment | 98.8% | 99.8% | 81.3% | N/A | N/A | N/A |

### Phylogenetic analysis of MAGs

To conduct phylogenetic analyses of the novel MAGs, genomes of *Coxiella*, *Rickettsia*, and *Midichloria* species were downloaded from The National Center for Biotechnology Information (NCBI) (Supplementary Table 4). For the *Rickettsiales* phylogeny, five outgroup species were included: *Orientia tsutsugamushi, Rickettsiales bacterium* Ac37b, *Anaplasma marginale*, *Escherichia coli* and a *Midichloriaceae* strain from *Ixodes persulcatus* (QG_MAG_2). For the *Coxiella* phylogeny, three outgroups were included: *Candidatus* Paracoxiella cheracis*, Rickettsiella massiliensis*, and *Rickettsiella* sp. isolate rickettsiella_Ird1 (Supplementary Table 4). From each genome, the 16S rRNA, 23S rRNA, and seven conserved single-copy marker genes (*atpA, dnaK, ftsZ, gltA, groEL, gyrB,* and *rpoB*) were extracted. Genes from *Coxiella* and *Rickettsiales* species were aligned separately using MAFFT, and the resulting alignments were concatenated. Maximum likelihood (ML) phylogenetic trees were constructed using IQ-TREE (v2.2.5) (59) with the parameters -m TEST -bb 1000 -alrt 1000 -nt AUTO. The best-fit substitution model selected by IQ-TREE was GTR+F+I+G4 for the *Rickettsiales* tree and TPM3+G4 for the *Coxiella* tree. Dot-plots were generated using D-Genies (v1.5.0) (60) to compare the MAGs with their closest common ancestors (Supplementary Figure 5).

### Pangenomic analysis of Midichloria species

Genomes from *Midichloria* spp. (n=31) were downloaded from NCBI and the European Nucleotide Archive (ENA) (Supplementary Table 5). These were combined with the *Midichloria* contigs generated in the current study. Taxonomic classification was determined with GTDB-Tk (v2.5.2) (56) with GTDB r226, with the classify (classify_wf) (Supplementary Table 6) and *de novo* workflows, placing the *Midichloria* genomes into a Rickettsiales phylogenetic tree with GTDB-Tk reference genomes (de_novo_wf --bacteria --taxa_filter o Rickettsiales --outgroup_taxon o Paracaedibacterales) (Supplementary Figure 5). Next, the average nucleotide identities (ANI) and amino acid identities (AAI) were compared between the *Midichloria* genomes using FastANI (v.1.34) (61) and CompareM (v0.1.2) (https://github.com/dparks1134/CompareM), respectively (Supplementary Table 5). MAGs that did not meet the AAI genus threshold (<65%) were excluded from the pangenome analysis. To avoid quasi-clonal representatives, *Midichloria* strains were dereplicated by setting an ANI threshold of 98%, and the representative with the highest genome completeness using CheckM2 (v1.0.2) (54) was selected (Supplementary Table 5). A pangenomic analysis of the remaining nine *Midichloria* spp. was performed with anvi’o (v8) (62) following the standard pangenomic workflow. Protein-coding genes were clustered with the default MCL inflation value of 2. Genes and metabolic pathways were annotated using the KEGG database with the anvi-run-kegg-kofams function, the Clusters of Orthologous Groups (COGs) database (COG20) using the anvi-run-ncbi-cogs function and the eggNOG database (v.5) by running eggNOG-mapper (v2.1.12) (63) on the genes and importing the results into anvi’o with the anvi-import-functions function (last accessed 12/11/2025). The concatenated *Midichloria* spp. single-copy marker genes from the *Rickettsiales* phylogenetic tree were extracted to create a *Midichloria*-only ML phylogenetic tree with IQ-TREE (v2.2.2.5) (59) (-m TEST -bb 1000 -alrt 1000 -nt AUTO). The best-fit model selected was GTR+F+G4. This phylogenetic tree was subsequently imported into anvi’o using the anvi-import-misc-data function to organise the genomes according to their evolutionary relationships.

## RESULTS

### Genome assemblies of novel *Coxiella, Rickettsia* and *Midichloria* species

Through short-read sequencing and the *de novo* metagenome assembly pipeline, a total of six complete or partial metagenome-assembled genomes (MAGs) were obtained (Table 2). The two complete and circularised genomes were from the *Haemaphysalis longicornis Coxiella*-like endosymbiont and the *Ixodes holocyclus Midichloria mitochondrii* Ixholo1. They shared high identity with previously sequenced MAGs, including a *Coxiella*-like endosymbiont from a *Haemaphysalis longicornis* sequenced in China (98.8% identity, GenBank: GCF_024730665.1) and another *Ixodes holocyclus Midichloria mitochondrii* isolate (“IholoSidney”, 99.8% identity; GenBank: GCA_030060845.1), respectively. The remaining four partial MAGs, *Midichloria mitochondrii* Ixholo2, a *Bothriocroton hydrosauri Coxiella* sp., a *Bothriocroton concolor Coxiella* sp., and a *Bothriocroton concolor Rickettsia* sp., do not appear to have been sequenced previously. The genome completeness of these strains/species ranges from 49.22% in the *Bothriocroton concolor* Rickettsia sp. to 71.5% in *Midichloria mitochondrii* Ixholo2 (Table 2).

### Phylogenetic analysis of novel *Coxiella, Rickettsia* and *Midichloria* MAGs

Maximum-likelihood (ML) phylogenetic analyses were conducted using 16S rRNA, 23S rRNA, and seven conserved single-copy marker genes to elucidate the evolutionary relationships between newly assembled MAGs and previously characterised *Coxiella* and *Rickettsiales* genomes. The ML tree of *Rickettsiales* revealed that the novel *Rickettsia* endosymbiont of *B. concolor* clusters within the non-pathogenic Bellii clade. It is highly similar (ANI of 99.0%) to *Rickettsia* sp. GXaj.114_MAG_1 of the tick *Amblyomma javanense* from China (GenBank: SAMEA118347715) (Supplementary Figure 5). These two strains form a sister group to the *Rickettsia* endosymbiont of the true bug, *Gonocerus acuteangulatus* (GenBank: OZ032147) (Figure 1A).

**Figure 1.**
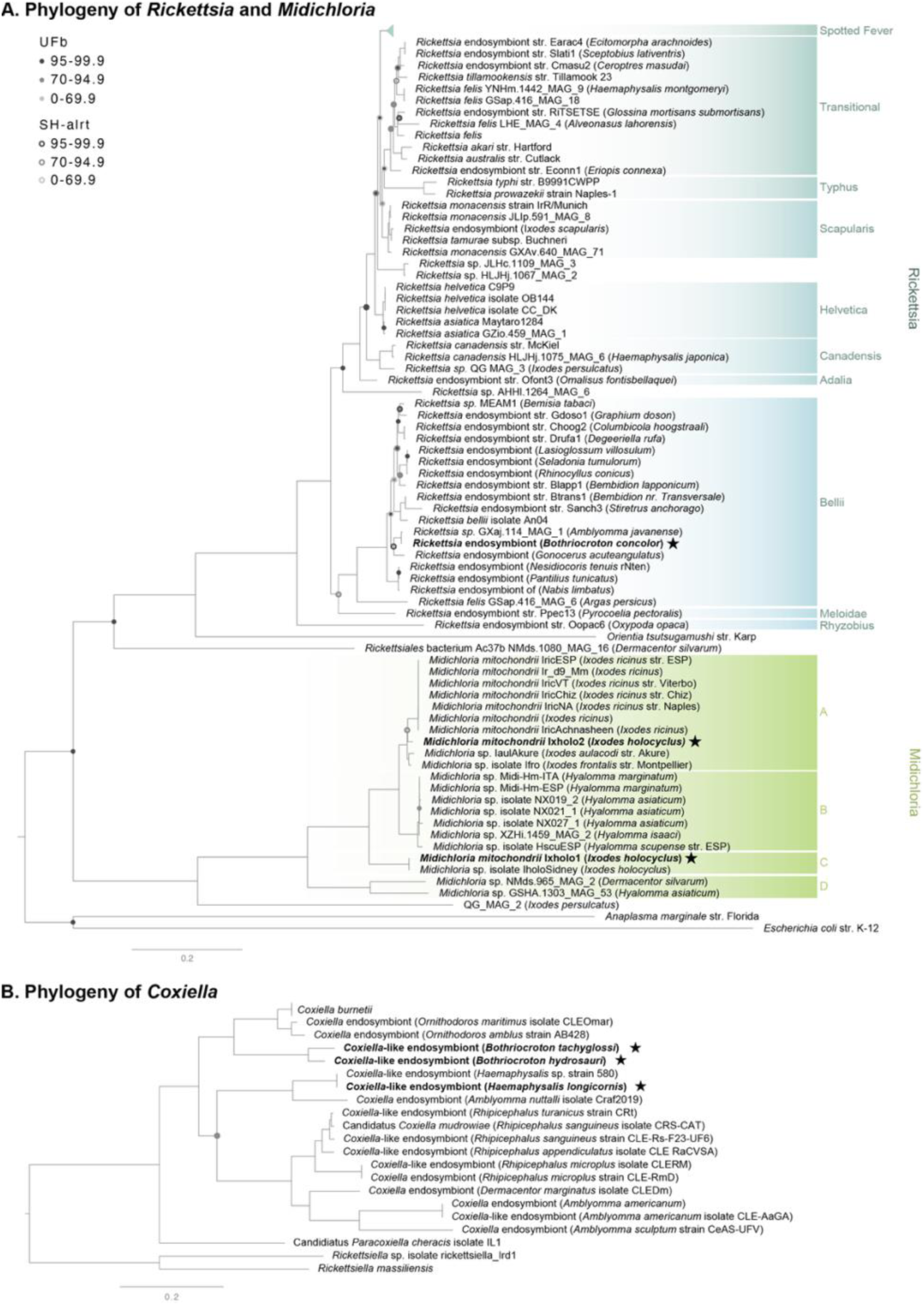
Phylogeny of *Rickettsiales* and *Coxiella* MAGs constructed using maximum Likelihood (ML) estimations based on concatenated 16S rRNA, 23S rRNA and single-copy marker genes (*atpA*, *dnaK*, *ftsZ*, *gltA*, *groEL*, *gyrB* and *rpoB*). **A.** The novel *Rickettsiales* MAGs, *Rickettsia* endosymbiont of *Bothricroton concolor*, *M. mitochondrii* Ixholo1 and *M. mitochondrii* Ixholo2 and **B.** the novel *Coxiella*-like endosymbionts of *B. concolor, B. hydrosauri* and *H. longicornis* are shown in bold, with a star. Bootstrap support values (<100) are reported on corresponding branches as circles. The inside circle shows the Ultra-fast bootstrap support value, and the circle outline shows the Shimodaira–Hasegawa approximate likelihood ratio test (SH-alrt) value. The scale-bar represents the number of expected changes per site. Sequence metadata, including GenBank accession numbers, are in Supplementary Table 4.

The *Rickettsiales* phylogenetic analyses additionally delineated four distinct clades within *Midichloria* (clades A, B, C and D) (Figure 1A). The *M. mitochondrii* ixholo1 MAG from this study forms a well-supported monophyletic clade (clade C) with a previously sequenced Ixholo1 genome (iholoSidney, GenBank: GCA_030060845.1) (UFBoot = 100; SH-aLRT = 100) (Figure 1A). In contrast, *M. mitochondrii* Ixholo2 clusters within the broader *Ixodes* clade (clade A), as sister to *M. mitochondrii* from European *Ixodes ricinus* ticks (UFBoot = 35.5; SH-aLRT = 89) (Figure 1A) (Supplementary Table 4).

The *Coxiella* ML phylogeny indicates that the *Coxiella*-like endosymbionts (CLEs) from *B. concolor* and *B. hydrosauri* form a distinct clade, which is sister to the clade comprising CLEs from *Ornithodoros* spp. and *Coxiella burnetii* (Figure 1B). Additionally, the CLE from *Haemaphysalis longicornis* is phylogenetically indistinguishable from a CLE previously sequenced from an unknown *Haemaphysalis* species in China (GenBank: CP084737), and together they form a sister group to the CLE from the tick *Amblyomma nuttalli* (GenBank: CP064834) (Figure 1B) (Supplementary Table 4).

### Genes involved in bacteria-tick symbiosis

Genes associated with bacteria-tick symbiosis were identified in the genomes of *Coxiella, Rickettsia* and *Midichloria* species (Figure 2). These include genes involved in the biosynthesis of heme and B vitamins, including biotin (B_7_), folate (B_9_), riboflavin (B_2_) (Supplementary Figure 6). The complete CLE genomes, including the novel CLE of *H. longicornis*, all encode the full biotin, folate, and riboflavin biosynthesis pathways, but only *gltX* from the heme pathway. The remaining two partially sequenced CLE MAGs of *B. concolor* and *B. hydrosauri* similarly encode genes of the three B vitamins but not the heme biosynthesis pathway (Figure 2).

**Figure 2.**
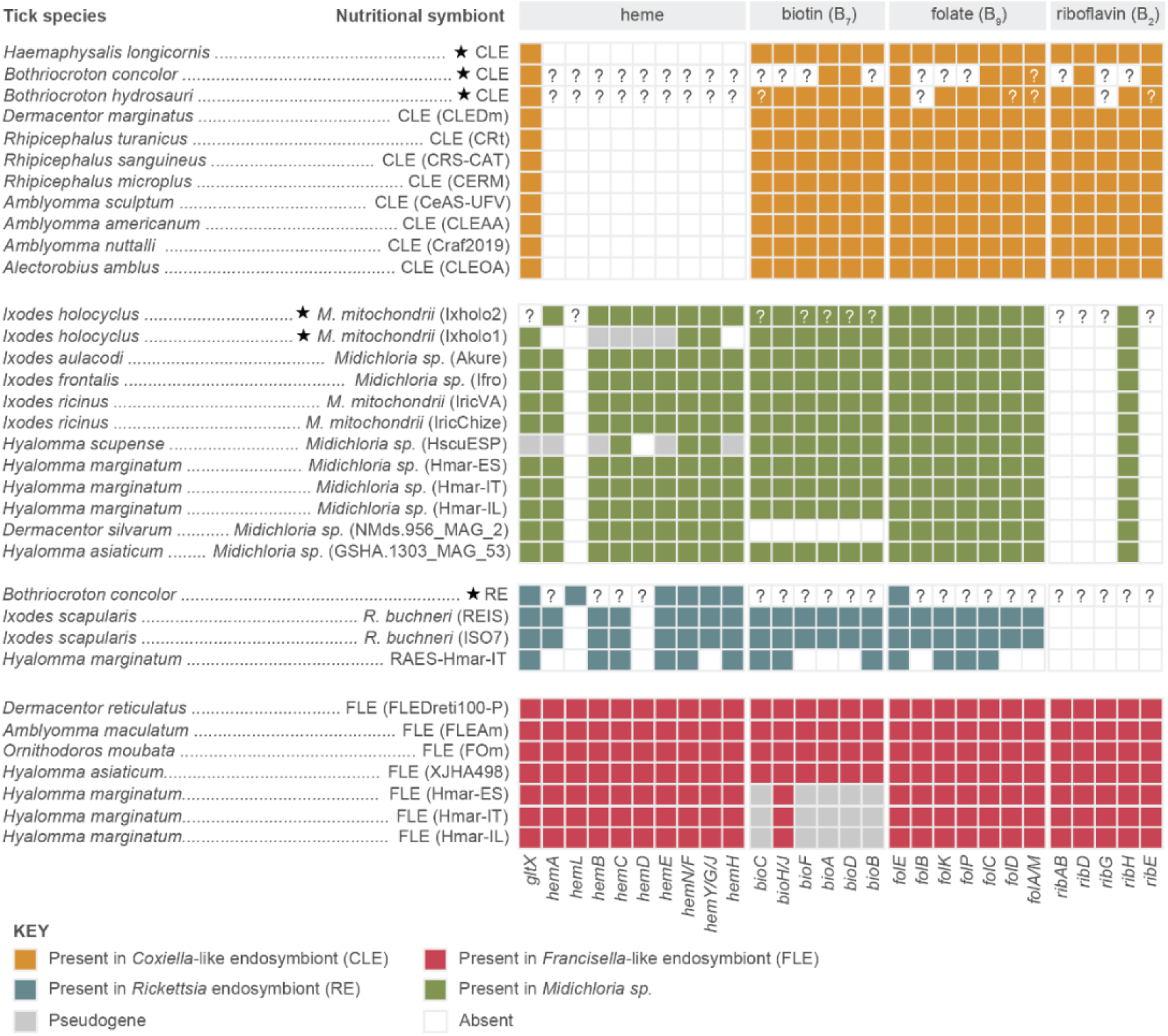
Genes involved in bacteria-tick endosymbiosis. Absence (white), presence (coloured) or pseudogenisation (grey) of genes from heme, biotin, folate, and riboflavin biosynthesis pathways. The tick host species are shown with their respective nutritional symbionts: *Coxiella*-like endosymbiont (CLE) (orange), *Rickettsia* endosymbiont (RE) (blue), *Francisella*-like endosymbiont (red) and *Midichloria mitochondrii* (green). The bacterial strain names are shown in brackets. For symbionts with partial genomes, partially recovered genes are represented with a white question mark, and genes not recovered are represented with a back question mark. Figure adapted from (12, 15).

All *M. mitochondrii* strains encode complete biotin and folate biosynthesis pathways, but only *ribH* of the riboflavin pathway, except for the *Midichloria sp.* NMds.956_MAG_2, which lacks the biotin operon. They additionally possess all the necessary genes for heme biosynthesis, except for *M. mitochondrii* Ixholo1 and *Midichloria* sp. HscuESP, which possess only three of the nine intact heme biosynthesis genes found in other strains, with the remaining six either absent or pseudogenised (Figure 2). In the partial genome of *Rickettsia* endosymbiont of *B. concolor* (Figure 2), no biotin and riboflavin genes, and only some heme (*gltX* and *hemLEFJH*) and folate (*folE*) biosynthesis genes were identified. No homologs of *cif* genes were detected.

### Pangenomic analysis of *Midichloria* species

To perform a *Midichloria* pangenomic analysis, we first confirmed the generic placement of the 31 *Midichloria* MAGs using AAI and GTDB-Tk. The AAI between all the MAGs were above 65% except *Midichloria* sp. QG_MAG_2, which shared 53% AAI with the other MAGs (Supplementary Table 5). GTDB-Tk classified all the MAGs as *Midichloria*, except for *Midichloria* sp. QG_MAG_2, NMds.965_MAG_2 and GSHA.1303_MAG_53 (Supplementary Table 6, Supplementary Figure 7). The *Midichloria* MAGs within the A, B and C clades shared <95% ANI with genomes within their clades, whereas those from clade D, *Midichloria* sp. NMds.965_MAG_2 and GSHA.1303_MAG_53 shared 80.97% ANI. The MAGs were then dereplicated for the pangenomic analysis by setting a 98% ANI threshold, resulting in nine *Midichloria* representatives (Figure 3).

**Figure 3.**
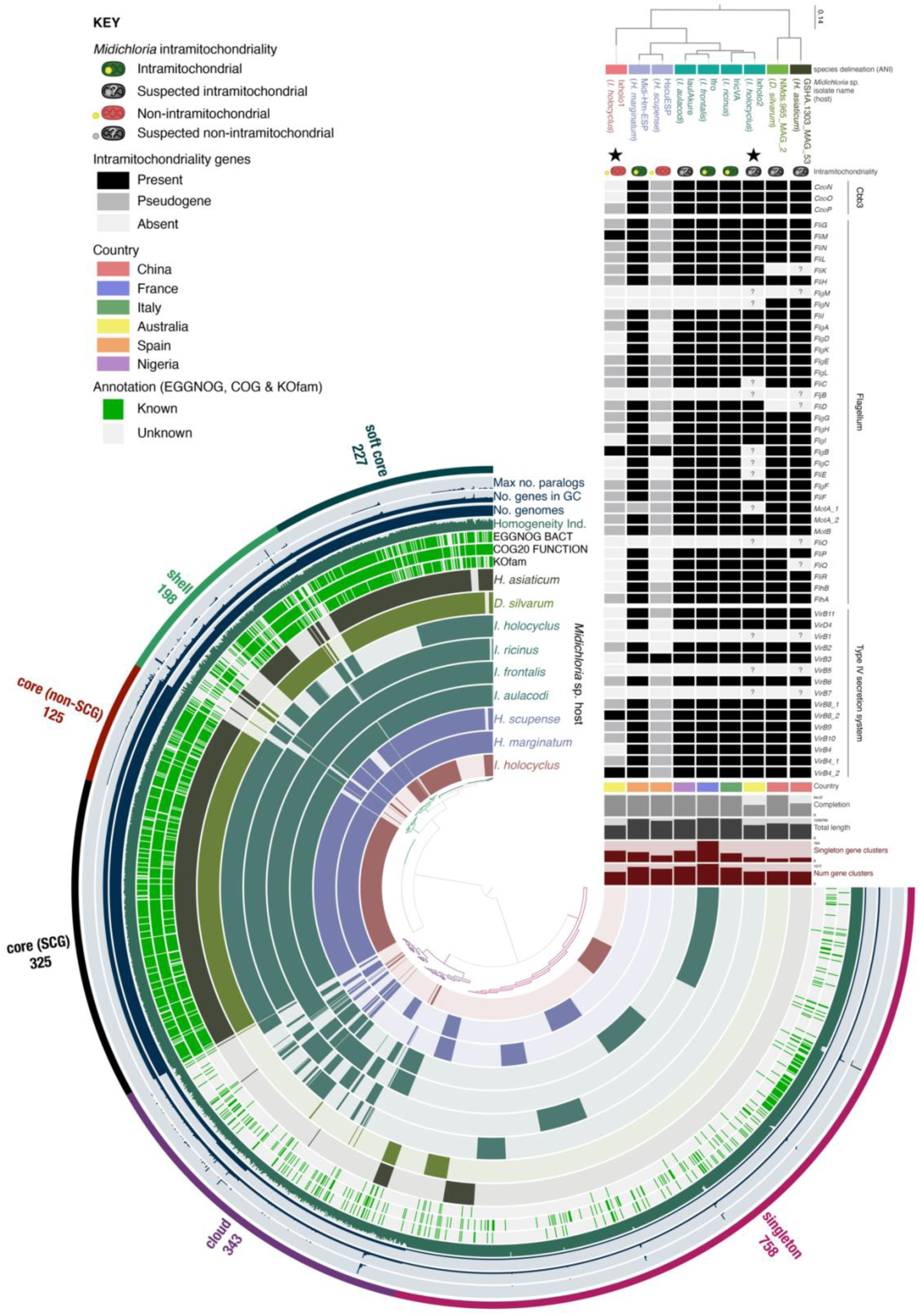
Pangenome analysis of *Midichloria* species. Pangenome representation of nine *Midichloria* MAGs, comprising 1,976 gene clusters with 9,700 genes, generated using the anvi’o software. Layers represent individual genomes, organised and coloured (brown, green, blue, purple and red) according to ANI values (>95% identity) and clustering in the *Midichloria* sp. maximum likelihood (ML) phylogeny reconstructed from concatenated 16s rRNA, 23s rRNA and seven single-copy marker genes (*atpA*, *dnaK*, *ftsZ*, *gltA*, *groEL*, *gyrB* and *rpoB*). The scale bar represents the number of substitutions per site. In the layers, dark colours indicate the presence of a gene cluster and light colour its absence. Core (single copy genes, and non-single copy genes), soft core, shell, cloud and singleton genes are labelled, with their respective number of gene clusters. Functional annotations of orthologs are shown above the genomes, including Kegg Orthology (KOfam), Clusters of Orthologous Genes (COG20 FUNCTION) and EggNOG-mapper (EGGNOG BACT). Genes with known annotations are green, and those with unknown annotations are off-white. The outer rings show the combined homogeneity of the gene clusters’ amino acid sequences, the number of contributing genomes, the number of genes in the gene clusters and the maximum number of paralogs. The intramitochondrial *Midichloria* strains are indicated by green mitochondria harbouring a bacterium, while the non-intramitochondrial *Midichloria* strains are represented by a bacterium outside a red mitochondrion. The hypothesised intramitochondriality of *Midichloria* strain of unknown localisation are shown in black and white. The presence (black), absence (white) or pseudogenisation (grey) of genes suspected to be involved in intramitochondriality are shown for each genome, including genes involved in the type IV secretion system, the flagellar genes and the cbb3 pathway (21, 64). For *Midichloria* strains with partial genomes, genes not recovered are represented with a question mark. The countries from which the ticks were sampled (China, France, Italy, Australia, Spain, and Nigeria) are colour-coded. Histograms represent the total number of gene clusters, number of singleton gene clusters, the total MAG length (ranging from 812 kilobases to 1.22 megabases) and percent (%) completion of the MAGs.

In these nine genomes, the anvi’o pangenomic analysis described 1,976 gene clusters with 9,700 genes. Of these gene clusters, 450 were core genes (including 325 single-copy genes (SCG) and 125 non-SCGs), 227 were soft core genes, 198 were shell genes, 343 cloud genes and 758 were singletons (Figure3). The number of gene clusters in the *Midichloria* spp. ranged from 791 in *M. mitochondrii* Ixholo1 to 1,277 in *M. mitochondrii* strain Ifro. The number of singleton gene clusters ranged from 33 in *Midichloria* sp. NMds.965_MAG_2 to 194 in *M. mitochondrii* strain Ifro, and were mostly annotated as hypothetical proteins in all nine MAGs. In *M. mitochondrii* Ixholo2, of the three cytochrome *cbb3*, 15 type IV secretion system (T4SS) and 34 flagellum genes, 40 were present, 12 were absent, and none were pseudogenised. This contrasts with *M. mitochondrii* Ixholo1, which had four present, 22 absent and 26 pseudogenised (Figure 3).

## DISCUSSION

### Australian ticks harbour non-pathogenic endosymbionts that are highly specific to tick genera

Shotgun metagenomic sequencing was conducted on four Australian tick species, *B. concolor, B. hydrosauri, H. longicornis* and *I. holocyclus*. The metagenomic analyses revealed that the ticks had diverse microbial communities (Supplementary Figure 4), and enabled the assembly of six MAGs. Of these six MAGs, two are circularised novel isolates from previously sequenced endosymbionts: CLE of *H. longicornis* and *M. mitochondrii* Ixholo1, and four are partial genomes of newly sequenced tick endosymbionts: CLE of *B. hydrosauri,* CLE of *B. concolor*, RE of *B. concolor* and *M. mitochondrii* Ixholo2. Past 16S rRNA gene sequencing of *H. longicornis* detected high abundances of *Coxiella* endosymbionts, suggesting that this CLE is abundant in *H. longicornis* around the world (65–67). The three CLEs lack the virulence genes encoding the Dot/Icm secretion system, and the RE of *B. concolor* clusters with *Rickettsia* species in the non-pathogenic Bellii clade. The absence of recognised virulence determinants and phylogenetic placement among known endosymbionts are consistent with an endosymbiotic rather than a recognised pathogenic lifestyle. Similar to the report of ticks examined from diverse locations (15), the Australian ticks in this study exhibit a high prevalence of one or two bacterial symbionts.

### Ticks require vitamin supplementation through nutritional symbionts

Ticks depend on symbionts for B vitamin provisioning as vertebrate blood is nutrient-poor. These B vitamins include riboflavin (vitamin B_2_), biotin (vitamin B_7_), and folate (vitamin B_9_), and have been experimentally proven to be essential for tick growth, reproduction and survival (68–72). The genes necessary for *de novo* synthesis of biotin (*bioABCFH/J*) and folate (*folEBKPCDA/M*) have been identified in all CLE and *Midichloria* sp. genomes previously analysed, as well as most FLE (Figure 2). All the biotin and folate biosynthesis genes were present in the CLE of *H. longicornis*, while only some were identified in the partial CLE of *B. concolor* and CLE of *B. hydrosauri* MAGs (Figure 2). The absence of some of these genes is likely due to incomplete sequencing. In *Midichloria* and *Rickettsia*, the biotin pathway is a compact operon (known as BOOM - Biotin synthesis Operon of Obligate intracellular Microbes) (73) which was acquired following an ancient lateral gene transfer event with another intracellular bacterium (15). This operon can drive the evolution toward nutritional symbiosis in intracellular bacteria that originally lacked the ability to synthesise biotin (15). Its absence in *Midichloria* sp. QG_MAG_2, *M. mitochondrii* NMds.965_MAG_2 and the Rickettsia MAG from *B. concolor* suggest they are facultative symbionts and not nutritional symbionts (74) (Figure 2). In contrast, the presence of the operon in *M. mitochondrii* GSHA.1303_MAG_53 and *M. mitochondrii* from the clade A, B and C lineages, indicates that the lateral gene transfer event may have taken place in a common ancestor of *M. mitochondrii* but was lost in *M. mitochondrii* NMds.965_MAG_2. Overall, the presence of biotin and folate biosynthesis genes in the CLE and *M. mitochondrii* MAGs demonstrate genomic capacity for nutritional symbiosis.

### Heme is likely to be synthesised by *M. mitochondrii* Ixholo2 and *Rickettsia* endosymbionts, but not *Coxiella*-like endosymbionts

Heme is an essential molecule required for biological processes including cellular respiration and binding of diatomic gases (75). Unlike most eukaryotes, ticks have lost most of the genes required for the synthesis and degradation of heme (76, 77). Since heme is essential for growth and reproduction, ticks rely on exogenous sources, such as blood, or production by nutritional symbionts. In most bacteria, heme is synthesised using the C5 pathway, which uses *gltX* and *hemL* in the first two steps (75) (Supplementary Figure 6). However, α-proteobacteria, such as *Rickettsia* and *Midichloria*, use the C4 (or Shemin) pathway, which uses *hemA* instead of *hemL* and *gltX* (75) (Supplementary Figure 6).

All the genes for the Shemin heme synthesis pathway were present in *M. mitochondrii* Ixholo2, suggesting that it is responsible for heme synthesis in *I. holocyclus* (Figure 2). In contrast, *M. mitochondrii* Ixholo1 appears to have lost several heme synthesis genes either completely or through pseudogenisation. This is similar to the loss of these genes in *Midichloria sp.* isolate HscuESP from *Hyalomma scupense* (Figure 2). Electron microscopy of *I. holocyclus* and *H. scupense* ovarian tissues indicate that both Ixholo1 and HscuESP lack intramitochondrial tropism (i.e. they do not colonise the host mitochondria) (64). Heme synthesis in *Midichloria* species may therefore be linked to intramitochondrial tropism.

The genome of the RE from *B. concolor* was largely incomplete. Nevertheless, it was shown to encode most of the genes required for heme biosynthesis (*gltX* and *hemLEFJH*) and therefore likely also has the genomic capacity to synthesise heme. The presence of the *hemL* and *gltX* genes in *B. concolor* RE suggests heme is synthesised through the C5 pathway, which is unusual for an α-proteobacterium. In contrast, the CLEs of *H. longicornis, B. concolor*, and *B. hydrosauri* do not encode any heme synthesis genes and therefore do not appear to synthesise heme, as has been observed in other CLEs (12). These tick species may acquire heme uniquely by salvaging it from the host during blood meals, and not from a nutritional symbiont. Overall, the results suggest that heme is synthesised by *M. mitochondrii* Ixholo2 and RE of *B. concolor,* but not *M. mitochondrii* xholo1, CLE of *H. longicornis, B. concolor*, and *B. hydrosauri*.

### Influence of tick endosymbionts on feeding, development and vector competence

In addition to providing essential nutrients and cofactors, endosymbionts can also influence tick feeding. For example, antibiotic-induced depletion of *Coxiella* endosymbionts in *Rhipicephalus sanguineus, Rhipicephalus microplus*, and *H. longicornis*, as well as *Francisella* endosymbionts in *Ornithodoros moubata* significantly compromised or stopped tick feeding and further moulting (69–71, 78). Zhong et. al (2021) experimentally demonstrated that *Coxiella* endosymbionts in *H. longicornis* regulate blood intake through the biosynthesis of chorismate, an important precursor for the biosynthesis of 5-hydroxytryptamine (5-HT; serotonin) (79). Genomic analysis of the MAGs revealed the presence of all the chorismate biosynthesis genes (*aroGBOEKAC*) in the Australian CLE of *H. longicornis*, four (*aroGEAC*) in the partial CLE of *H. hydrosayri* MAG, one (*aroG*) in the partial CLE of *B. conocolor* MAG, and one (*aroK*) in *M. mitochondrii* Ixholo2. These results suggest that the CLEs sequenced in this study may similarly influence blood feeding through the regulation of serotonin biosynthesis. In Australia, *Haemaphysalis longicornis* is a vector of *Theileria orientalis,* which causes Oriental theileriosis in bovine and was estimated to cost $20 million per year nationally in a 2015 report (80, 81). Similarly, *Bothriocroton hydrosauri* is an important vector of *Rickettsia honei*, the aetiological agent of Flinders Island spotted fever in humans (19). Ticks treated with the herbicide glyphosate, which inhibits chorismate biosynthesis, have been experimentally shown to reduce feeding and present an interesting alternative approach for tick control (79).

The importance of endosymbionts on tick development has additionally been demonstrated experimentally through the rearing of tetracycline-treated ticks. This treatment resulted in reduced levels of bacteria in the progeny of *R. microplus*, resulting in an inhibition of development beyond the metanymph stage (71). Similarly, larvae of *M. mitochondrii*-free *I. ricinus* performed poorly during blood feeding, highlighting the influence of *M. mitochondrii* on tick development (82). To determine if the MAGs sequenced in this study are derived from obligate endosymbionts, similar experimental studies should be conducted on Australian ticks. The importance of nutritional symbionts on tick physiology and the taxon-specific nature of tick endosymbionts could allow for taxon-specific biocontrols of ticks through anti-microbiota vaccines.

Endosymbionts can also influence tick vector competence by either promoting or inhibiting pathogens (83, 84). For example, endosymbionts can inhibit pathogen proliferation through antibiotic activity, as well as inhibit colonisation and survival through resource competition (83–87). Whether the endosymbionts from which MAGs were sequenced in this study have a positive or negative influence on pathogen colonisation and survival remains unknown and requires further investigation. Complex endosymbiont-pathogen interactions pose an interesting opportunity to leverage pathogen inhibition and decrease tick vector competence.

### Characterising tick endosymbionts is important for the prevention of misidentified pathogens by diagnostic tools

*Coxiella*-like and *Rickettsia*-like endosymbionts belong to the same genera as Australian tick-borne pathogens. These pathogens include *Coxiella burnetii*, the aetiological agent of Q fever, *Rickettsia honei*, the causative agent of Flinders Island spotted fever (88), *Rickettsia australis*, the etiological agent of Queensland tick typhus (89), and *R. honei* subsp. marmionii, the causative agent of Australian spotted fever (90). Due to the sequence similarities between bacteria of the same genus, it is important to target pathogen-specific regions of DNA when creating PCR-based diagnostic tools to avoid false positive results. For example, *C. burnetii* prevalence used to rely on the amplification of the *IS111* gene, including in Australia (91–93). However, this gene was later characterised in CLE throughout Africa, Europe and South America (94), demonstrating that *IS111* is not *C. burnetii-*specific and that Q fever detection assays based solely on this gene misidentified CLE. Since this discovery, molecular detection of *C. burnetii* uses additional marker genes such as *groEL* and *com1* (95). Sequencing symbiont genomes therefore ensures that diagnostic tools accurately detect pathogens and overcome false positives.

### Average nucleotide and amino acid identities suggest the *Midichloria* genus could be divided into five species

*Rickettsiales* phylogenetic reconstruction based on 16S rRNA, 23S rRNA, and seven single-copy marker genes revealed that the genus *Midichloria* is split into four clades, however, the number of species is unclear. The phylogeny was congruent with the previously reported topology for Ixholo1 and Ixholo2 derived from 16S rRNA analysis, placing Ixholo2 in the *I. ricinus M. mitochondrii* (IricVA) clade (clade A) and Ixholo1 in its own clade (Figure 1) (20). Previous 16S rRNA and genomic ANI analyses on *Midichloria* additionally suggest the genus is split into three species (96, 97). It is generally accepted that an ANI and AAI ≥ 95% between two MAGs indicates they are of the same species, and < 95% indicates a different species (98–100). By these standards, the *M. mitochondrii* clades A, B and C represent three separate species, as previously shown by Melis et. al 2025, and the *Midichloria sp.* NMds.965_MAG_2 and *Midichloria sp.* GSHA.1303_MAG_53 from clade D are two separate species (Supplementary Table 5) (97). Furthermore, an AAI of 65-95% between two MAGs indicates they are of the same genus, while an AAI below 65% indicates a different genus (98). Clade D *Midichloria* spp. (NMds.965_MAG_2 and GSHA.1303_MAG_53) share between 70.24 and 72.37% AAI with the other *Midichloria* species, suggesting they are *Midichloria* species. However, GTDB-Tk did not classify them as *Midichloria*, leaving the taxonomic placement of these two species unclear (Supplementary Tables 5 and 6). Lastly, *Midichloria* sp. QG MAG_2 (SAMEA118348234) shares ∼53% AAI identity with the remaining *Midichloria* species. This falls below the accepted cutoff of 65 to 95%, suggesting that this MAG is of a different genus and has been wrongly annotated as *Midichloria*. To formally describe a new uncultured species, a MAG is required to have a genome completeness of >80% (with <5% contamination), genetic discreteness (ANI/AAI standards), phylogenetic placement, bioinformatics-based functional and phenotypic predictions and microscopic identification (98). Further analyses are therefore required for reclassification of the *Midichloria* genus, particularly microscopic identification by FISH or other visualisation techniques for species outside clade A.

### Intramitochondriality of *M. mitochondrii* Ixholo2

*Ixodes holocyclus* harbours the two *M. mitochondrii* strains, Ixholo1 and Ixholo2, however, the exact role of the two strains remains unclear. Following a blood meal, *M. mitochondrii* from *I. ricinus* substantially increase in numbers, suggesting they may be involved in the moulting process (101). Genomic and metatranscriptomics analyses additionally suggest that *M. mitochondrii* Ixholo1 aid in osmotic regulation, stress response and nutrition through biotin and folate synthesis for their host (21, 22). Amplicon sequencing additionally uncovered a third *M. mitochondrii* strain, which differs from the Ixholo2 V1 to V3 region of the 16S rRNA gene by one nucleotide (Supplementary Figure 6). This strain was found in two of the five samples, a fed adult female (sample 1) and an unfed adult male (sample 4) (Supplementary Figure 4). Both these samples had high *M. mitochondrii* Ixholo2 loads, suggesting there may be a positive correlation between the two strains.

The genomic data of Ixholo2 and *Midichloria* pangenomic analyses revealed genomic signatures of mitochondrial tropism, including the cytochrome cbb3, heme, T4SS, flagellum genes and genes involved in membrane and cell wall structure, biosynthesis, and structure (Figure 3, Supplementary Figure 2). The cytochrome cbb3 complex is an alternative version of the terminal oxidase of the electron transport chain in Proteobacteria. This complex has a higher affinity for oxygen, making it important for colonising microaerobic environments such as the mitochondria (21, 102). An intramitochondrial symbiont such as Ixholo2 could therefore salvage oxygen from mitochondria to produce ATP to either be used by the symbiont or exported and used by the host. *M. mitochondrii* Ixholo2 additionally have intact *de novo* heme synthesis pathways, similarly to other intramitochondrial *M. mitochondrii*. This may be because Fe^2+^ is essential for the final step in the heme synthesis pathway, and mitochondria are iron-rich, unlike most environments (21, 103). Furthermore, because heme is energetically expensive to synthesise, intramitochondriality of Ixholo2 would presumably allow heme to be seamlessly synthesised within the mitochondria, allowing tighter control by the host without losing intermediate products. The T4SS and flagellum are suggested to be used for adhesion and invasion of host cells and mitochondria, as well as transporting effector molecules such as DNA and proteins (64). The loss of these genes, as well as genes involved in membrane and cell wall structure, biosynthesis, and structure in *M. mitochondrii* Ixholo1 and *M. mitochondrii* HscuE indicates a simplification of the external cell layers (Supplementary Figure 8) (64).

Through the analysis of 95 *I. holocyclus* transcriptomic datasets, a recent study found variation in Ixholo1 and Ixholo2 frequencies in different sexes and life stages (21). They found that Ixholo2 transcripts were more abundant in adult males (74.4%), while Ixholo1 transcripts were more abundant in nymphs and adult females (93.7% and 72.5%, respectively), suggesting specialisation of the symbiont for different tick sexes and life stages. Since adult males *I. holocyclus* do not feed, Ixholo2 may therefore be supplementing heme, B vitamins and ATP for their survival until they mate with a female.

Electron microscopy of ovarian cells of four unengorged adult female *I. holocyclus* found no evidence for mitochondrial tropism of symbiotic bacteria (20). Since Ixholo2 appears to be most abundant in males, future studies should therefore test for intramitochondriality in different sexes, as well as life stages and tissue types of *I. holocyclus*. The retention of the cytochrome *cbb3*, heme, T4SS, flagellum, cell wall and membrane genes in *M. mitochondrii* Ixholo2 is suggestive of the endosymbiont exhibiting mitochondrial tropism.

## CONCLUSION

Shotgun metagenomic sequencing was performed on four Australian tick species, including *B. concolor*, *B. hydrosauri* and *H. longicornis* which were sequenced for the first time. Through bacterial enrichment using filtration, six bacterial MAGs were retrieved. The absence of virulence factors and presence of B vitamin and heme biosynthesis genes in the five CLE and *Midichloria* MAGs is indicative of nutritional mutualism which is essential for tick hematophagy. The CLEs additionally harbour genes of the shikimate pathway, which modulates blood feeding in ticks by regulating serotonin biosynthesis. The retention of the cytochrome *cbb3*, heme, T4SS, flagellum, cell wall and membrane genes in *M. mitochondrii* Ixholo2 also suggests that the endosymbiont exhibits mitochondrial tropism. Tick microbiomes are dominated by non-pathogenic microbes, which are often overshadowed by pathogens. These include the endosymbionts, which can influence pathogen transmission, are important for the development of diagnostic tools and could allow for taxon-specific biocontrols of ticks through anti-microbiota vaccines, highlighting the need for further research in this field

## Author statements

### Author contributions

Laurene Leclerc: Conceptualization, Formal analysis, Funding acquisition, Investigation, Methodology, Validation, Visualization, Writing – original draft

Xabier Vázquez-Campos: Formal analysis, Methodology, Writing – review & editing

Julia Meltzer: Formal analysis, Methodology, Writing – review & editing

Olivier Duron: Conceptualization, Methodology, Writing – review & editing

Julien Amoros: Formal analysis, Methodology, Writing – review & editing

Brendan P. Burns: Conceptualization, Methodology, Resources, Supervision, Writing – review & editing

Nathan Lo: Conceptualization, Methodology, Resources, Supervision, Writing – review & editing

## Conflicts of interest

*The author(s) declare that there are no conflicts of interest*.

## Funding information

This work was funded by The Karl McManus Foundation. This work was supported by “Investissements d’Avenir” managed by the Agence Nationale de la Recherche (ANR, France, ref. ANR-25-CE02-7068, to OD and Laboratoire d’Excellence CEBA, ref. ANR-10-LABX-25-01 to OD).

## Ethical approval

N/A

## Consent for publication

N/A

## Acknowledgements

This research includes computations using the computational cluster Katana supported by Research Technology Services at UNSW Sydney (https://doi.org/10.26190/669x-a286). We would also like to acknowledge Dr Ann Mitrovic, the Koala Conservation Australia, the Byron Bay Wildlife Hospital, and the Wandanian Wallaby & Kangaroo Rehabilitation Centre for generously donating ticks.

## Notes

https://www.ncbi.nlm.nih.gov/bioproject/PRJNA1457570

### Competing Interest Statement

The authors have declared no competing interest.

## References

1. Rochlin I, Toledo A. Emerging tick-borne pathogens of public health importance: a mini-review. Journal of Medical Microbiology. 2020;69(6):781.

2. Bruley M, Duron O. Multi-locus sequence analysis unveils a novel genus of filarial nematodes associated with ticks in French Guiana. Parasite. 2024;31:14.

3. Binetruy F, Duron O. Molecular detection of Cercopithifilaria, Cruorifilaria and Dipetalonema-like filarial nematodes in ticks of French Guiana. Parasite. 2023;30:24.

4. Bezerra-Santos MA, de Macedo LO, Nguyen V-L, Manoj RR, Laidoudi Y, Latrofa MS, et al. Cercopithifilaria spp. in ticks of companion animals from Asia: new putative hosts and vectors. Ticks and Tick-Borne Diseases. 2022;13(4):101957.

5. Moutailler S, Popovici I, Devillers E, Vayssier-Taussat M, Eloit M. Diversity of viruses in *Ixodes ricinus*, and characterization of a neurotropic strain of Eyach virus. New Microbes and New Infections. 2016;11:71–81.

6. Du L-F, Shi W, Cui X-M, Fan H, Jiang J-F, Bian C, et al. Genome-resolved metagenomics reveals microbiome diversity across 48 tick species. Nature Microbiology. 2025;10(10):2631–45.

7. Buysse M, Koual R, Binetruy F, de Thoisy B, Baudrimont X, Garnier S, et al. Detection of Anaplasma and Ehrlichia bacteria in humans, wildlife, and ticks in the Amazon rainforest. Nature Communications. 2024;15(1):3988.

8. Ni X-B, Cui X-M, Liu J-Y, Ye R-Z, Wu Y-Q, Jiang J-F, et al. Metavirome of 31 tick species provides a compendium of 1,801 RNA virus genomes. Nature microbiology. 2023;8(1):162–73.

9. Duron O, Koual R, Musset L, Buysse M, Lambert Y, Jaulhac B, et al. Novel chronic anaplasmosis in splenectomized patient, Amazon rainforest. Emerging Infectious Diseases. 2022;28(8):1673.

10. Hollenhorst C. Factors of Climate Change and Their Influence on Lyme Disease Infection Rates: Harvard University; 2024.

11. Hussain S, Perveen N, Hussain A, Song B, Aziz MU, Zeb J, et al. The symbiotic continuum within ticks: Opportunities for disease control. Frontiers in Microbiology. 2022;13:854803.

12. Duron O. Nutritional symbiosis in ticks: singularities of the genus *Ixodes*. Trends in Parasitology. 2024;40(8):696–706.

13. Buysse M, Duron O. Evidence that microbes identified as tick-borne pathogens are nutritional endosymbionts. Cell. 2021;184(9):2259–60.

14. Duron O, Gottlieb Y. Convergence of nutritional symbioses in obligate blood feeders. Trends in Parasitology. 2020;36(10):816–25.

15. Buysse M, Floriano AM, Gottlieb Y, Nardi T, Comandatore F, Olivieri E, et al. A dual endosymbiosis supports nutritional adaptation to hematophagy in the invasive tick *Hyalomma marginatum*. Elife. 2021;10:e72747.

16. Bonnet SI, Binetruy F, Hernández-Jarguín AM, Duron O. The tick microbiome: why non-pathogenic microorganisms matter in tick biology and pathogen transmission. Frontiers in Cellular and Infection Microbiology. 2017;7:236.

17. Floriano AM, El-Filali A, Amoros J, Buysse M, Jourdan-Pineau H, Sprong H, et al. Comparative genomics of Rickettsiella bacteria reveal variable metabolic pathways potentially involved in symbiotic interactions with arthropods. Peer Community Journal. 2025;5.

18. Amoros J, Buysse M, Floriano AM, Moumen B, Vavre F, Bouchon D, et al. Diversity and spread of cytoplasmic incompatibility genes among maternally inherited symbionts. PLoS Genetics. 2025;21(9):e1011856.

19. Barker SC, Barker D. Ticks of Australasia: 125 species of ticks in and around Australia. Zootaxa. 2023;5253(1):1–670.

20. Beninati T, Riegler M, Vilcins I-ME, Sacchi L, McFadyen R, Krockenberger M, et al. Absence of the symbiont *Candidatus* Midichloria mitochondrii in the mitochondria of the tick *Ixodes holocyclus*. FEMS Microbiology Letters. 2009;299(2):241–7.

21. Leclerc L, Mattick J, Burns BP, Sassera D, Hotopp JD, Lo N. Metatranscriptomics provide insights into the role of the symbiont *Midichloria mitochondrii* in *Ixodes ticks*. FEMS Microbiology Ecology. 2024;100(12):fiae133.

22. Olivieri E, Epis S, Castelli M, Boccazzi IV, Romeo C, Desirò A, et al. Tissue tropism and metabolic pathways of *Midichloria mitochondrii* suggest tissue-specific functions in the symbiosis with *Ixodes ricinus*. Ticks and Tick-borne Diseases. 2019;10(5):1070–7.

23. Greay TL, Evasco KL, Evans ML, Oskam CL, Magni PA, Ryan UM, et al. Illuminating the bacterial microbiome of Australian ticks with 16S and *Rickettsia*-specific next-generation sequencing. Current Research in Parasitology & Vector-Borne Diseases. 2021;1:100037.

24. Gofton AW, Doggett S, Ratchford A, Oskam CL, Paparini A, Ryan U, et al. Bacterial profiling reveals novel “*Ca.* Neoehrlichia”, *Ehrlichia*, and *Anaplasma* species in Australian human-biting ticks. PLoS One. 2015;10(12):e0145449.

25. Egan SL, Loh S-M, Banks PB, Gillett A, Ahlstrom L, Ryan UM, et al. Bacterial community profiling highlights complex diversity and novel organisms in wildlife ticks. Ticks and Tick-borne Diseases. 2020;11(3):101407.

26. Chandra S, Harvey E, Emery D, Holmes EC, Šlapeta J. Unbiased characterization of the microbiome and virome of questing ticks. Frontiers in Microbiology. 2021;12:627327.

27. Egan SL, Taylor CL, Banks PB, Northover AS, Ahlstrom LA, Ryan UM, et al. The bacterial biome of ticks and their wildlife hosts at the urban–wildland interface. Microbial Genomics. 2021;7(12).

28. Egan SL, Taylor CL, Austen JM, Banks PB, Northover AS, Ahlstrom LA, et al. Haemoprotozoan surveillance in peri-urban native and introduced wildlife from Australia. Current Research in Parasitology & Vector-Borne Diseases. 2021;1:100052.

29. Gofton AW, Doggett S, Ratchford A, Ryan U, Irwin P. Phylogenetic characterisation of two novel *Anaplasmataceae* from Australian *Ixodes holocyclus* ticks: ‘*Candidatus* Neoehrlichia australis’ and ‘*Candidatus* Neoehrlichia arcana’. International Journal of Systematic and Evolutionary Microbiology. 2016;66(10):4256–61.

30. Greay TL, Gofton AW, Paparini A, Ryan UM, Oskam CL, Irwin PJ. Recent insights into the tick microbiome gained through next-generation sequencing. Parasites & Vectors. 2018;11(1):1–14.

31. Hussain-Yusuf H, Stenos J, Vincent G, Shima A, Abell S, Preece ND, et al. Screening for *Rickettsia*, *Coxiella* and *Borrelia* species in ticks from Queensland, Australia. Pathogens. 2020;9(12):1016.

32. Loh S-M, Gofton AW, Lo N, Gillett A, Ryan UM, Irwin PJ, et al. Novel *Borrelia* species detected in echidna ticks, *Bothriocroton concolor*, in Australia. Parasites & Vectors. 2016;9:1–7.

33. Chalada MJ, Stenos J, Vincent G, Barker D, Bradbury RS. A molecular survey of tick-borne pathogens from ticks collected in central Queensland, Australia. Vector-Borne and Zoonotic Diseases. 2018;18(3):151–63.

34. Pace NR, Stahl DA, Lane DJ, Olsen GJ. The analysis of natural microbial populations by ribosomal RNA sequences. Advances in Microbial Ecology: Springer; 1986. p. 1–55.

35. Olsen GJ, Lane DJ, Giovannoni SJ, Pace NR, Stahl DA. Microbial ecology and evolution: a ribosomal RNA approach. Annual Reviews in Microbiology. 1986;40(1):337–65.

36. Harvey E, Rose K, Eden J-S, Lo N, Abeyasuriya T, Shi M, et al. Extensive diversity of RNA viruses in Australian ticks. Journal of Virology. 2019;93(3):10.1128/jvi.01358-18.

37. Gofton AW, Blasdell KR, Taylor C, Banks PB, Michie M, Roy-Dufresne E, et al. Metatranscriptomic profiling reveals diverse tick-borne bacteria, protozoans and viruses in ticks and wildlife from Australia. Transboundary and Emerging Diseases. 2022;69(5):e2389–e407.

38. Tyson GW, Chapman J, Hugenholtz P, Allen EE, Ram RJ, Richardson PM, et al. Community structure and metabolism through reconstruction of microbial genomes from the environment. Nature. 2004;428(6978):37–43.

39. Parry RH, Teo EJ, Petrone ME, Stewart A, Burnard D, Barker SC. Metagenomic resolution of spotted-fever group Rickettsia tasmanensis and novel DNA viruses in Australian wildlife ticks, with spatial modelling of Rickettsia exposure zones. bioRxiv. 2025:2025.07. 08.663691.

40. Bushnell B. BBMap: a fast, accurate, splice-aware aligner. 2014.

41. Andrews S. FastQC: a quality control tool for high throughput sequence data. Babraham Bioinformatics, Babraham Institute, Cambridge, United Kingdom; 2010.

42. Bankevich A, Nurk S, Antipov D, Gurevich AA, Dvorkin M, Kulikov AS, et al. SPAdes: a new genome assembly algorithm and its applications to single-cell sequencing. Journal of Computational Biology. 2012;19(5):455–77.

43. Nurk S, Meleshko D, Korobeynikov A, Pevzner PA. metaSPAdes: a new versatile metagenomic assembler. Genome Research. 2017;27(5):824–34.

44. Li D, Liu C-M, Luo R, Sadakane K, Lam T-W. MEGAHIT: an ultra-fast single-node solution for large and complex metagenomics assembly via succinct de Bruijn graph. Bioinformatics. 2015;31(10):1674–6.

45. Pronk LJ, Medema MH. Whokaryote: distinguishing eukaryotic and prokaryotic contigs in metagenomes based on gene structure. Microbial Genomics. 2022;8(5):000823.

46. Langmead B, Salzberg SL. Fast gapped-read alignment with Bowtie 2. Nature Methods. 2012;9(4):357–9.

47. Li H, Handsaker B, Wysoker A, Fennell T, Ruan J, Homer N, et al. The sequence alignment/map format and SAMtools. Bioinformatics. 2009;25(16):2078–9.

48. Sahlin K. Strobealign: flexible seed size enables ultra-fast and accurate read alignment. Genome Biology. 2022;23(1):260.

49. Steinegger M, Söding J. MMseqs2 enables sensitive protein sequence searching for the analysis of massive data sets. Nature Biotechnology. 2017;35(11):1026–8.

50. Kutuzova S, Piera P, Nor Nielsen K, Olsen NS, Riber L, Gobbi A, et al. Binning meets taxonomy: TaxVAMB improves metagenome binning using bi-modal variational autoencoder. bioRxiv. 2024:2024.10. 25.620172.

51. Wickham H. ggplot2. Wiley Interdisciplinary Reviews: Computational Statistics. 2011;3(2):180–5. DOI:.

52. Sievert C. Interactive web-based data visualization with R, plotly, and shiny: Chapman and Hall/CRC; 2020.

53. Allaire J. RStudio: integrated development environment for R. Boston, MA. 2012;770(394):165–71.

54. Parks DH, Imelfort M, Skennerton CT, Hugenholtz P, Tyson GW. CheckM: assessing the quality of microbial genomes recovered from isolates, single cells, and metagenomes. Genome Research. 2015;25(7):1043–55.

55. Seppey M, Manni M, Zdobnov EM. BUSCO: assessing genome assembly and annotation completeness. Gene Prediction: Methods and Protocols: Springer; 2019. p. 227–45.

56. Chaumeil P-A, Mussig AJ, Hugenholtz P, Parks DH. GTDB-Tk v2: memory friendly classification with the genome taxonomy database. Bioinformatics. 2022;38(23):5315–6.

57. Darling AC, Mau B, Blattner FR, Perna NT. Mauve: multiple alignment of conserved genomic sequence with rearrangements. Genome Research. 2004;14(7):1394–403.

58. Seemann T. Prokka: rapid prokaryotic genome annotation. Bioinformatics. 2014;30(14):2068–9.

59. Minh BQ, Schmidt HA, Chernomor O, Schrempf D, Woodhams MD, Von Haeseler A, et al. IQ-TREE 2: new models and efficient methods for phylogenetic inference in the genomic era. Molecular Biology and Evolution. 2020;37(5):1530–4.

60. Cabanettes F, Klopp C. D-GENIES: dot plot large genomes in an interactive, efficient and simple way. PeerJ. 2018;6:e4958.

61. Musiał K, Petruńko L, Gmiter D. Simple approach to bacterial genomes comparison based on Average Nucleotide Identity (ANI) using fastANI and ANIclustermap. Acta Universitatis Lodziensis Folia Biologica et Oecologica. 2024;18:66–71.

62. Eren AM, Esen ÖC, Quince C, Vineis JH, Morrison HG, Sogin ML, et al. Anvi’o: an advanced analysis and visualization platform for ‘omics data. PeerJ. 2015;3:e1319.

63. Cantalapiedra CP, Hernández-Plaza A, Letunic I, Bork P, Huerta-Cepas J. eggNOG-mapper v2: functional annotation, orthology assignments, and domain prediction at the metagenomic scale. Molecular Biology and Evolution. 2021;38(12):5825–9.

64. Floriano AM, Batisti Biffignandi G, Castelli M, Olivieri E, Clementi E, Comandatore F, et al. The evolution of intramitochondriality in Midichloria bacteria. Environmental Microbiology. 2023. 10.1111/1462-2920.16446.

65. Sang MK, Park JE, Song DK, Jeong JY, Hwang HJ, Kim HW, et al. Characterization of *Haemaphysalis longicornis* microbiome collected from different regions of Korean peninsula. Entomological Research. 2022;52(6):271–80.

66. Ponnusamy L, Travanty NV, Watson DW, Seagle SW, Boyce RM, Reiskind MH. Microbiome of invasive tick species *Haemaphysalis Longicornis* in North Carolina, USA. Insects. 2024;15(3):153.

67. Esteves E, Obellianne C, Garba A, Sarkar SL, Schuler MG, Hermance ME. Defining the kinetics of severe fever with thrombocytopenia syndrome virus acquisition and dissemination in naturally-infected *Haemaphysalis longicornis*. Frontiers in Cellular and Infection Microbiology. 2025;15:1706970.

68. Zhong J, Jasinskas A, Barbour AG. Antibiotic treatment of the tick vector *Amblyomma americanum* reduced reproductive fitness. PLoS One. 2007;2(5):e405.

69. Ben-Yosef M, Rot A, Mahagna M, Kapri E, Behar A, Gottlieb Y. *Coxiella*-like endosymbiont of *Rhipicephalus sanguineus* is required for physiological processes during ontogeny. Frontiers in Microbiology. 2020;11:493.

70. Duron O, Morel O, Noël V, Buysse M, Binetruy F, Lancelot R, et al. Tick-bacteria mutualism depends on B vitamin synthesis pathways. Current Biology. 2018;28(12):1896–902. e5.

71. Guizzo MG, Parizi LF, Nunes RD, Schama R, Albano RM, Tirloni L, et al. A *Coxiella* mutualist symbiont is essential to the development of *Rhipicephalus microplus*. Scientific Reports. 2017;7(1):17554.

72. Li L-H, Zhang Y, Zhu D. Effects of antibiotic treatment on the fecundity of *Rhipicephalus haemaphysaloides* ticks. Parasites & Vectors. 2018;11(1):242.

73. Driscoll TP, Verhoeve VI, Brockway C, Shrewsberry DL, Plumer M, Sevdalis SE, et al. Evolution of Wolbachia mutualism and reproductive parasitism: insight from two novel strains that co-infect cat fleas. PeerJ. 2020;8:e10646.

74. Hunter DJ, Torkelson JL, Bodnar J, Mortazavi B, Laurent T, Deason J, et al. The Rickettsia endosymbiont of Ixodes pacificus contains all the genes of de novo folate biosynthesis. PloS one. 2015;10(12):e0144552.

75. Kořený L, Oborník M, Horáková E, Waller RF, Lukeš J. The convoluted history of haem biosynthesis. Biological Reviews. 2022;97(1):141–62.

76. Jia N, Wang J, Shi W, Du L, Sun Y, Zhan W, et al. Large-scale comparative analyses of tick genomes elucidate their genetic diversity and vector capacities. Cell. 2020;182(5):1328–40. e13.

77. Perner J, Hajdusek O, Kopacek P. Independent somatic distribution of heme and iron in ticks. Current Opinion in Insect Science. 2022;51:100916.

78. Zhang C-M, Li N-X, Zhang T-T, Qiu Z-X, Li Y, Li L-W, et al. Endosymbiont CLS-HI plays a role in reproduction and development of Haemaphysalis longicornis. Experimental and Applied Acarology. 2017;73(3):429–38.

79. Zhong Z, Zhong T, Peng Y, Zhou X, Wang Z, Tang H, et al. Symbiont-regulated serotonin biosynthesis modulates tick feeding activity. Cell Host & Microbe. 2021;29(10):1545–57. e4.

80. Lakew BT, Eastwood S, Walkden-Brown SW. Epidemiology and transmission of *Theileria orientalis* in Australasia. Pathogens. 2023;12(10):1187.

81. Lane J, Jubb T, Shephard R, Webb-Ware J, Fordyce G. MLA Final Report: Priority list of endemic diseases for the red meat industries. Meat and Livestock Australia, Sydney, Australia. M a L Australia (Ed). 2015.

82. Guizzo MG, Hatalová T, Frantová H, Zurek L, Kopáček P, Perner J. *Ixodes ricinus* ticks have a functional association with *Midichloria mitochondrii*. Frontiers in Cellular and Infection Microbiology. 2023;12:1930.

83. Gall CA, Reif KE, Scoles GA, Mason KL, Mousel M, Noh SM, et al. The bacterial microbiome of *Dermacentor andersoni* ticks influences pathogen susceptibility. The ISME Journal. 2016;10(8):1846–55.

84. Narasimhan S, Rajeevan N, Liu L, Zhao YO, Heisig J, Pan J, et al. Gut microbiota of the tick vector *Ixodes scapularis* modulate colonization of the Lyme disease spirochete. Cell Host & Microbe. 2014;15(1):58–71.

85. Chang X, Li X, Pei Y, Deng E, Wu S, Jiang J, et al. Roles of Tick Symbiotic Microorganisms in Pathogen Transmission. Zoonoses. 2025;5(1):972.

86. Khogali R, Bastos A, Getange D, Bargul JL, Kalayou S, Ongeso N, et al. Exploring the microbiomes of camel ticks to infer vector competence: insights from tissue-level symbiont-pathogen relationships. Scientific Reports. 2025;15(1):5574.

87. Cull B, Burkhardt NY, Wang X-R, Thorpe CJ, Oliver JD, Kurtti TJ, et al. The *Ixodes scapularis* symbiont *Rickettsia buchneri* inhibits growth of pathogenic Rickettsiaceae in tick cells: implications for vector competence. Frontiers in Veterinary Science. 2022;8:748427.

88. Graves SR, Stewart L, Stenos J, Stewart RS, Schmidt E, Hudson S, et al. Spotted fever group rickettsial infection in south-eastern Australia: isolation of rickettsiae. Comparative Immunology, Microbiology and Infectious Diseases. 1993;16(3):223–33.

89. Andrew R, Bonnin J, Williams S. Tick typhus in north Queensland. 1946:253–8.

90. Unsworth NB, Stenos J, Graves SR, Faa AG, Cox GE, Dyer JR, et al. Flinders Island spotted fever rickettsioses caused by “marmionii” strain of *Rickettsia honei*, Eastern Australia. Emerging Infectious Diseases. 2007;13(4):566.

91. Banazis MJ, Bestall AS, Reid SA, Fenwick SG. A survey of Western Australian sheep, cattle and kangaroos to determine the prevalence of *Coxiella burnetii*. Veterinary Microbiology. 2010;143(2-4):337–45.

92. Bennett MD, Woolford L, Banazis MJ, O’Hara AJ, Warren KS, Nicholls PK, et al. *Coxiella burnetii* in western barred bandicoots (*Perameles bougainville*) from Bernier and Dorre Islands in Western Australia. EcoHealth. 2011;8(4):519–24.

93. Potter AS, Banazis MJ, Yang R, Reid SA, Fenwick SG. Prevalence of *Coxiella burnetii* in western grey kangaroos (*Macropus fuliginosus*) in Western Australia. Journal of Wildlife Diseases. 2011;47(4):821–8.

94. Duron O. The IS1111 insertion sequence used for detection of *Coxiella burnetii* is widespread in *Coxiella*-like endosymbionts of ticks. FEMS Microbiology Letters. 2015;362(17):fnv132.

95. Mathews KO, Phalen D, Sheehy PA, Herbert CA, Brandimarti ME, Conaty JR, et al. Molecular detection and characterisation of Coxiella burnetii in Australian native wildlife species. FEMS Microbiology Letters. 2025:fnaf060.

96. Buysse M, Duron O. Multi-locus phylogenetics of the *Midichloria* endosymbionts reveals variable specificity of association with ticks. Parasitology. 2018;145(14):1969–78.

97. Melis S, Gammuto L, Castelli M, Nardi T, Bisaglia B, Duron O, et al. Genetic and genomic variability of *Spiroplasma* and *Midichloria* endosymbionts associated with the tick *Ixodes frontalis*. ISME Communications. 2025:ycaf202.

98. Konstantinidis KT, Rosselló-Móra R, Amann R. Uncultivated microbes in need of their own taxonomy. The ISME Journal. 2017;11(11):2399–406.

99. Potapov AM, Beaulieu F, Birkhofer K, Bluhm SL, Degtyarev MI, Devetter M, et al. Feeding habits and multifunctional classification of soil-associated consumers from protists to vertebrates. Biological Reviews. 2022;97(3):1057–117.

100. Richter M, Rosselló-Móra R. Shifting the genomic gold standard for the prokaryotic species definition. Proceedings of the National Academy of Sciences. 2009;106(45):19126–31.

101. Sassera D, Lo N, Bouman EA, Epis S, Mortarino M, Bandi C. “*Candidatus* Midichloria” endosymbionts bloom after the blood meal of the host, the hard tick *Ixodes ricinus*. Applied and Environmental Microbiology. 2008;74(19):6138–40.

102. Pitcher RS, Watmough NJ. The bacterial cytochrome cbb3 oxidases. Biochimica et Biophysica Acta (BBA)-Bioenergetics. 2004;1655:388–99.

103. Ward DM, Cloonan SM. Mitochondrial iron in human health and disease. Annual Review of Physiology. 2019;81:453–82.

