## Supplementary Figures 1-8 for "Characterisation and genomic analysis of bacterial nutritional endosymbionts in Australian ticks from shotgun metagenomic sequencing"

#### Supplementary methods

##### 16S rRNA (V1-V3) amplicon sequencing

To validate our DNA extraction and bacterial enrichment protocols, five samples were subjected to 16S rRNA (V1-3) amplicon sequencing, which was performed on extracted DNA at the Ramaciotti Centre for Genomics on an Illumina MiSeq v3 2×300 base pairs (bp) sequencing run using PCR primers 27f and 519r (Supplementary Tables 1 and 2). Amplicon sequence data were analysed using the DADA2 algorithm (v.1.32.0) (29) in RStudio (v4.4.1) (30). Reads were filtered and trimmed (truncLen=c(260,260), maxN=0, maxEE=c(3,5), truncQ=2, rm.phix=TRUE, compress=TRUE, multithread=TRUE, trimLeft = c(20, 17)), denoised by merging paired reads (maxMismatch = 1, minOverlap = 10), and chimeras removed. Read taxonomic assignment was performed using the SILVA database (v138.1) (31), and visualised with ggplot2 (v3.5.2) (32) histograms (Supplementary Figures 1 and 2). To compare *M. mitochondrii spp*. abundance in the different libraries, 16S rDNA read counts of *M. mitochondrii* Ixholo1, Ixholo2 and a third *M. mitochondrii* ‘Ixholo3’ were extracted (Supplementary Table 1, Supplementary Figures 2 and 3).

### Supplementary Figures

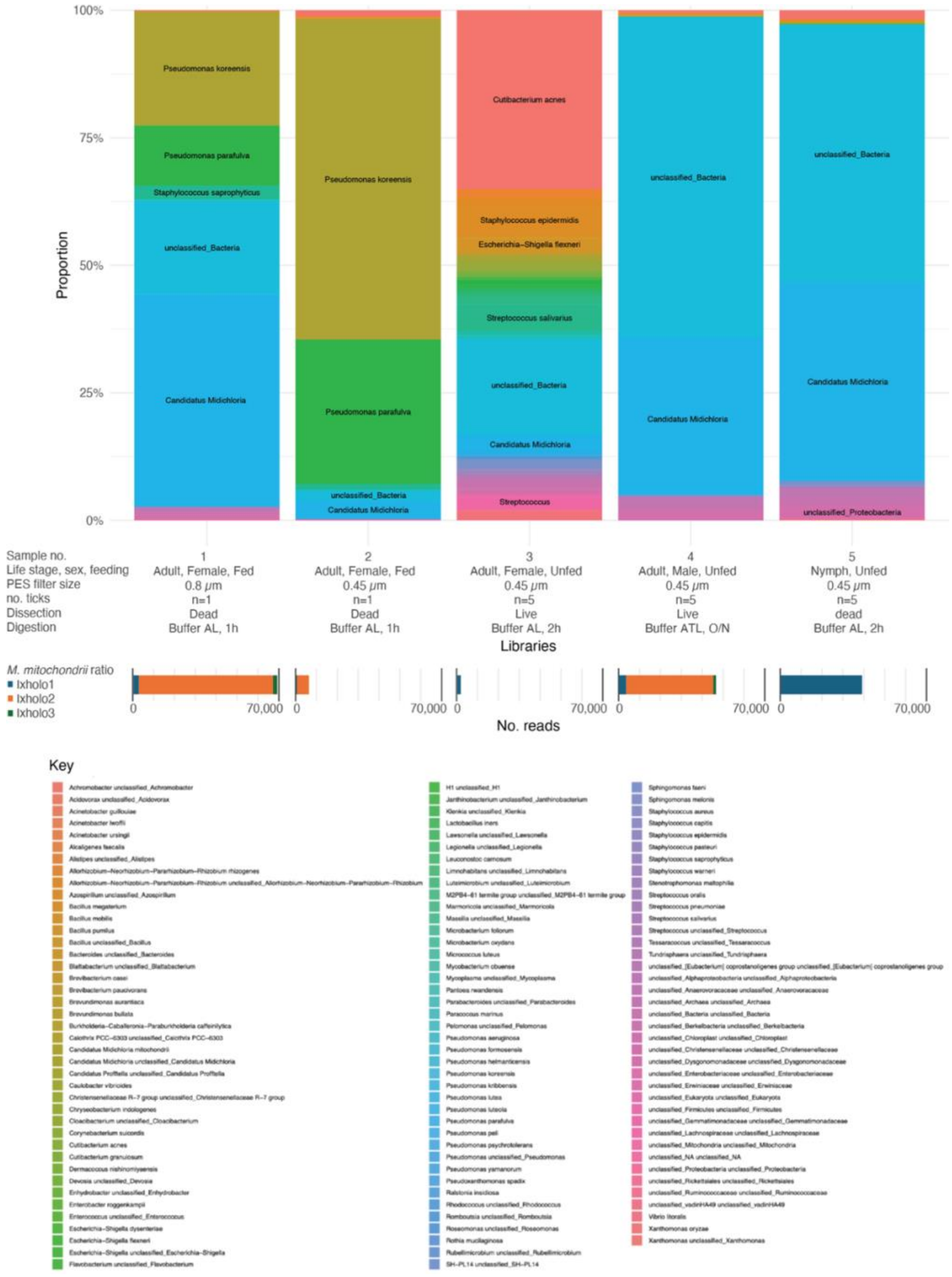


**Supplementary Figure 1. Proportion of bacterial genera (and species if known) in *I. holocyclus* 16S rRNA (V1-V3) amplicon sequencing libraries.** Information on the ticks in each library is shown below the respective columns, including tick life stage, sex, engorgement level, size of PES filter, number of ticks pooled and if they were dissected dead or alive. The number of *M. mitochondrii* Ixholo1 (ASV4, in blue), Ixholo2 (ASV1, in orange), and Ixholo3 (ASV13, in green) reads are also shown below the respective library columns.

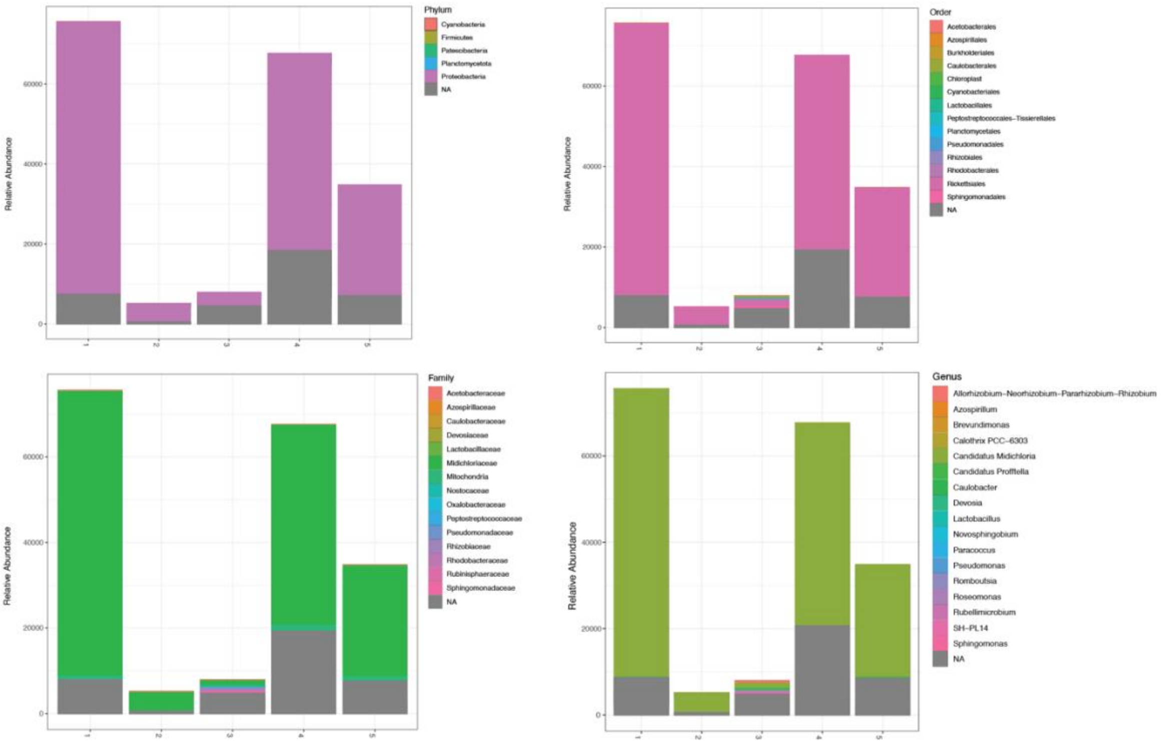


**Supplementary Figure 2.** Number of bacterial reads at different classification levels (Phylum, Order, Family and Genus) in *I. holocyclus* 16S rRNA (V1-V3) amplicon sequencing libraries.

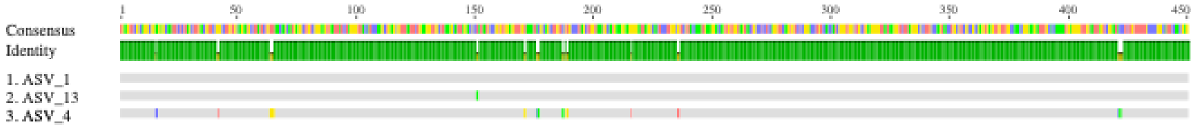


**Supplementary Figure 3. MAFFT alignment of *M. mitochondrii* Ixholo1 (ASV4), Ixholo2 (ASV1) and Ixholo3 (ASV13) 16S rRNA (V1-V3) using Geneious Prime.** Coloured lines in the alignment indicate Single-nucleotide polymorphism (SNPs). There is one SNP between ASV1 and ASV13 (99.8% identity), and 13 SNPs between ASV1 and ASV4 (97.1% identity). Consensus identity is shown above the alignment (green = 100%, yellow = 33.3%). Nucleotide length is shown above the alignment.

*See attached file, Supplementary_Figure_4.html*

**Supplementary Figure 4. Histogram showing the bacterial composition of each library based on assembled contigs.** Contigs were taxonomically classified to the species level using MMseqs2, and relative abundance was estimated from read coverage using Strobealign. Only contigs taxonomically assigned at the genus level are included, and unassigned contigs are ignored. These profiles should be interpreted with caution, as assembly-based abundance estimates are influenced by several known biases, including sequencing, assembly, binning, taxonomic annotation, and genome size biases (e.g., organisms with small genomes such as *M. mitochondrii* may be underrepresented). As a result, this histogram reflects contig-level representation rather than a quantitative reconstruction of the true microbial community within the ticks. Histogram was plotted using ggplot2 and Plotly.

A. *Midichloria* sp. Ixholo1 versus *Midichloria* sp. isolate IholoSidney (GCA_030060845.1)

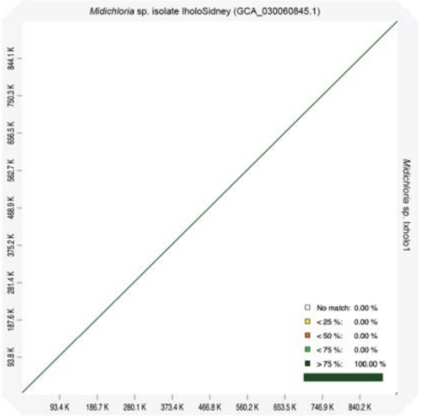


B. *Midichloria* sp. Ixholo2 versus *Midichloria mitochondrii* IricVA (CP002130)

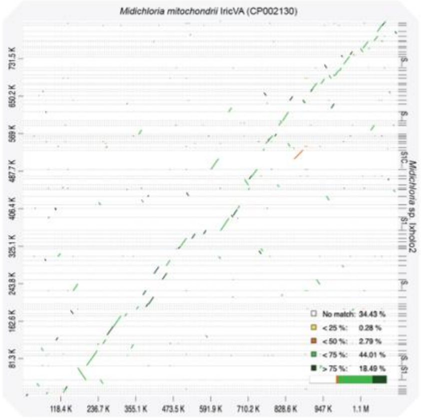


C. CLE of *H. longicornis* versus CLE of *Haemaphysalis* sp. (CP084737)

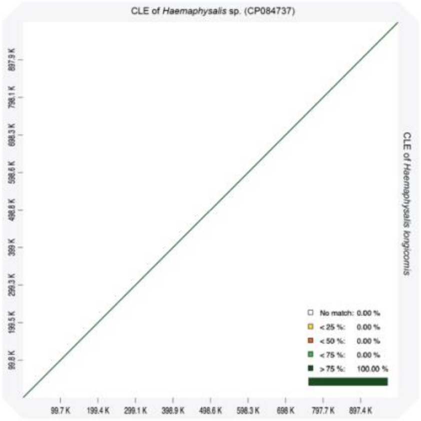


D. CLE of *Bothriocroton concolor* versus CLE of *Bothriocroton hydrosauri*

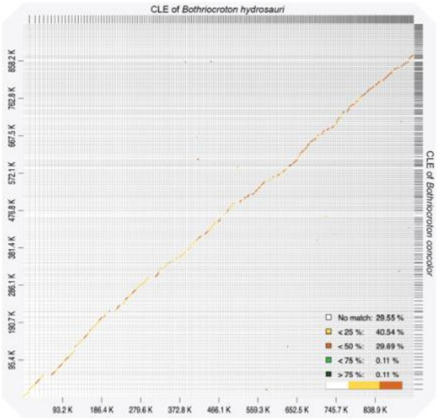


E. *Rickettisa* sp. of *B. concolor* versus *Rickettsia sp.* of *Amblyomma javanense* (SAMEA118347715)

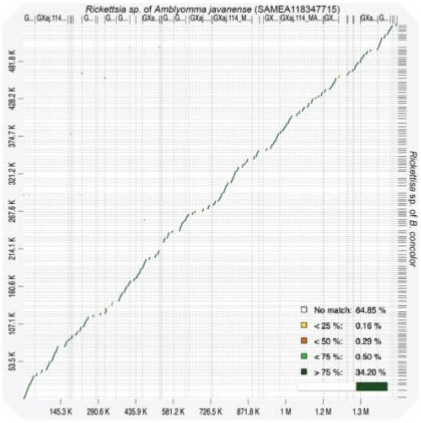


**Supplementary Figure 5. D-Genies plots of MAGs characterised in this study and their closest common ancestor, determined from the phylogenetic analysis (see main text).** GenBank accession numbers are shown in parentheses. The contig pairwise identities are colour-coded, with the percentage of MAG corresponding to each pairwise identity range (see key).

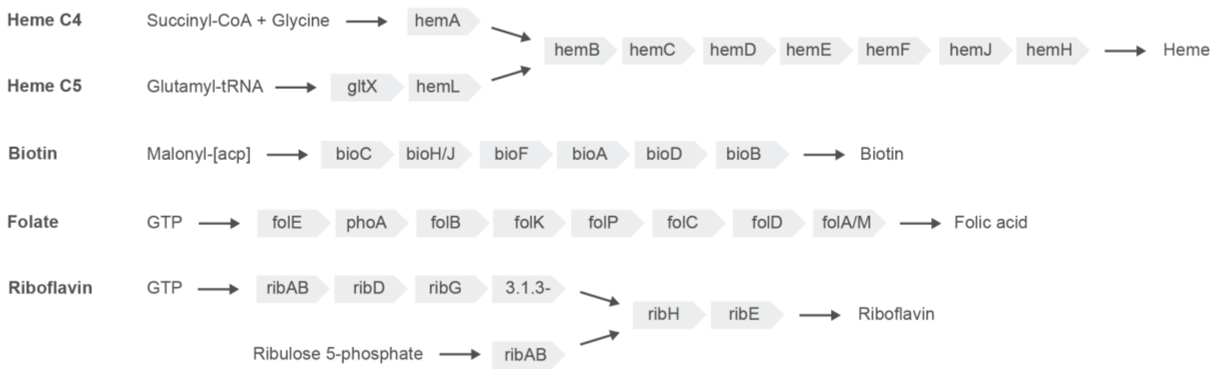


**Supplementary Figure 6.** Heme (C4 or Shemin and C5) and B-vitamins (biotin, folate and riboflavin) biosynthetic pathways of tick symbionts.

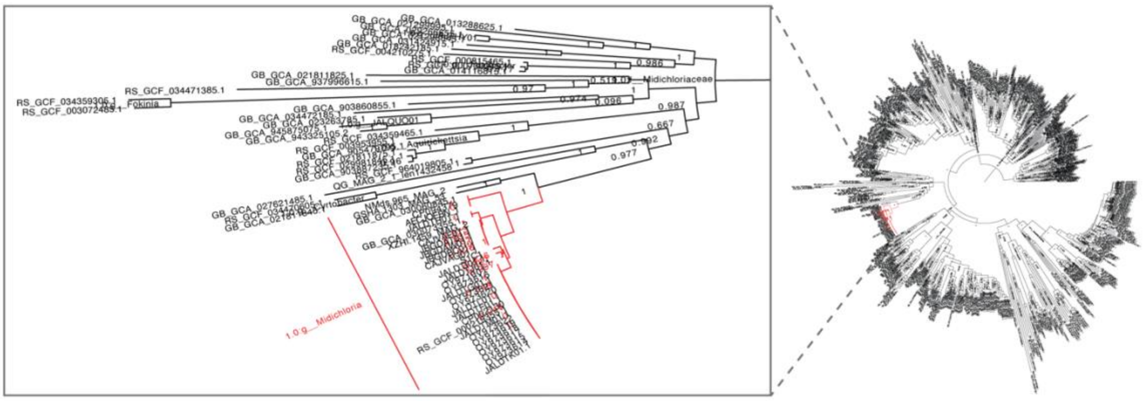

**Supplementary Figure 7.** GTDB-Tk tree of *Rickettsiales* with zoom into *Midichloriaseae*. Branches representing *Midichloria* species are coloured red. Leaves represent accession numbers.

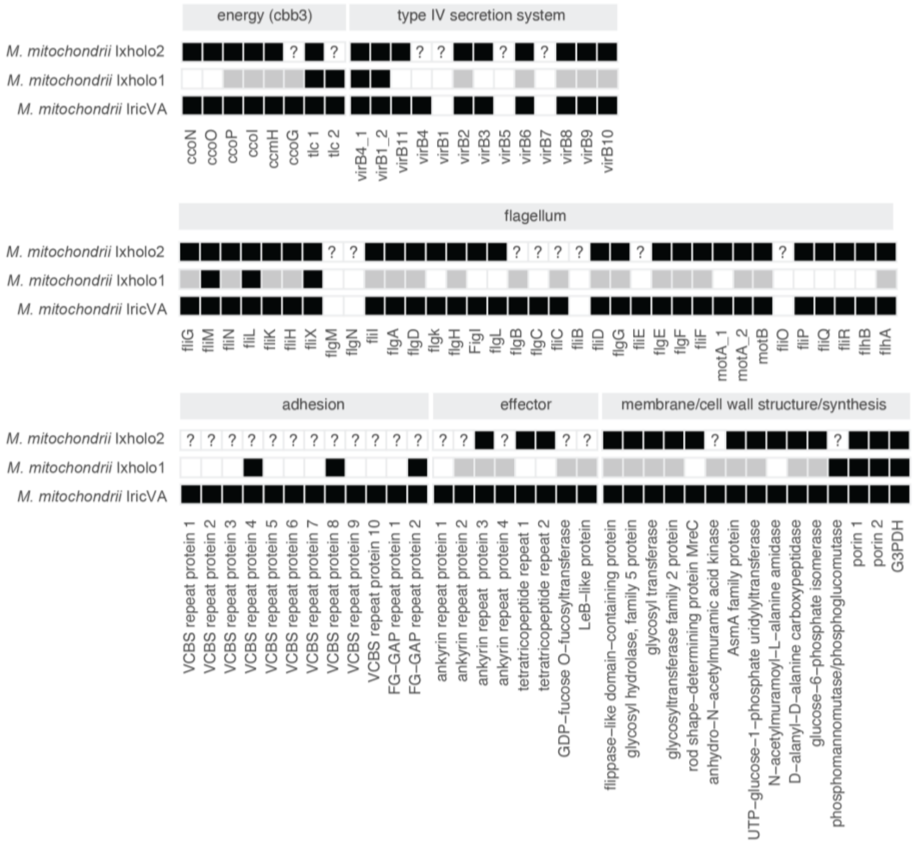


**Supplementary Figure 8.** Genes involved in *M. mitochondrii* intramitochondriality. Absence (white), presence (black) or pseudogenisation (grey; fragmented or truncated) of genes suggested to be involved in *M. mitochondrii* intramitochondriality. Including the two Australian *I. holocyclus M. mitochondrii* strains (Ixholo1 and Ixholo2) and the European *I. ricinus M. mitochondrii,* IricVA. The intramitochondriality genes include the type IV secretion system, flagellum, adhesion and effector proteins, membrane and cell wall structure synthesis genes and genes involved in the *cbb3* pathway. For symbionts with partial genomes, genes that were partially recovered are represented with a white question mark, and genes not recovered are represented with a back question mark.
